## Supplementary Material for "Action does not enhance but attenuates predicted touch" (from another version of this preprint: version 3, 2023-06-30)

**Just Noticeable Difference (JND) analysis**

From the psychophysical fits, we extracted the just noticeable difference (JND), which represents the participants’ force discrimination capacity between conditions. In Experiment 1, the JNDs did not significantly differ between any of the conditions (all p values > 0.05). In Experiment 2, the JNDs were not significantly different between conditions (*p value*s > 0.05), except in the *contact* condition, where the JNDs were significantly higher than in the *baseline* condition (*t*(29) = -2.52, *p* = 0.017, *d* = -0.46, *CI^95^ =* [-0.06 -0.01]). Finally, in Experiment 3, the JNDs were significantly lower in the *baseline* condition than in the *contact* condition (*t*(29) = -3.61, *p* <.001, *d* = -0.66, *CI^95^* = [-0.09 -0.02]) and the *no-contact* condition (*t*(29) = -2.77, *p* = 0.010, *d* = -0.51, *CI^95^* = [-0.08 -0.01]), but there were no significant differences in the JNDs between the *contact* and *no-contact* conditions (*t*(29) = -0.65, *p* = 0.521, *d* = -0.12, *CI^95^* = [-0.02 0.04], *BF*_01_ = 4.233). Thus, in contrast to the PSEs, there were no significant differences in the JNDs between the *contact* and *no-contact* conditions in any of the three experiments, demonstrating that the two conditions yielded the same discrimination capacity.

**Kinematic analysis**

To ensure that the movements in both the *contact* and *no-contact* conditions were comparable, participants were trained to make the same movement with their right index finger between two visual markers positioned 5 cm apart in Experiments 1 and 2. This training was not required for Experiment 3, in which participants were blindfolded and unaware that the distance sensor had been retracted on *no-contact* trials.

Given that the right index finger movement ends on impact with the force sensor in the *contact* condition but not in the *no-contact* condition, the kinematics of the finger movements are likely to differ between conditions. The active finger was slightly closer to the passive finger at the time of the *test* force in the *contact* compared to the *no-contact* condition in Experiments 1 and 2 (by less than 5 mm). Specifically, in Experiment 1, the position of the right index finger at the time of the *test* force was marginally closer to the left index finger in the *contact* condition than in the *no-contact* condition (mean difference = 2.32 mm, s.e.m. = 1.14: *t*(29) = -2.04, *p* = 0.051, *d* = 0.37, *CI^95^* = [-0.74 0.01], *BF*_01_ = 0.845. In Experiment 2, the position of the right index finger at the time of the *test* force was slightly closer to the left index finger in the *contact* condition than in the *no-contact* condition (mean difference = 4 mm, s.e.m. = 1.8: *t*(29) = -2.29, *p* = 0.030, *d* = 0.42, *CI^95^* = [-0.79 -0.04]. In Experiment 3, participants were unaware that they would miss the force sensor in the *no-contact* trials, so with no force sensor to stop the movement, their right index finger continued further downwards. Here, the position of the right index finger at the time of the *test* force was closer to the left index finger in the *no-contact* condition than in the *contact* condition (mean difference = 25.7 mm, s.e.m. = 2.0: *t*(29) = 12.79, *p* < .001, *d* = 2.34, *CI^95^* = [1.63 3.03]). Critically, the closer position of the right index finger to the left index finger in the *no-contact* trials compared to the *contact* trials cannot explain the attenuation findings in the *no-contact* condition since the attenuation in the *contact* trials was actually stronger compared to that in the *no-contact* trials (**Main Text**).

It could be argued that our differences between the *contact* and *no-contact* conditions across experiments are driven by differences in the timing of the two setups. This explanation is unlikely since we observed robust attenuation in the *no-contact* condition of Experiment 3 but not in Experiments 1 and 2 while the setups remained the same. To further control that the attenuation observed in the *no-contact* condition of Experiment 3 but not in the *no-contact* conditions of Experiments 1 and 2 was not driven by timing differences in the participants’ movement between experiments, we calculated the time the *test* force was delivered in the *contact* and *no-contact* conditions with respect to their movement end point (*i.e.*, the force sensor press in the *contact* condition and the lowest vertical position in the *no-contact* condition). On average, the *test* force was applied 24 ms earlier in the *no-contact* compared to the *contact* condition in Experiment 1 (s.e.m. = 4.20), 11 ms earlier in the *no-contact* compared to the *contact* condition in Experiment 2 (s.e.m. = 4.53) and 15 ms earlier in the *no-contact* compared to the *contact* condition in Experiment 3 (s.e.m. = 2.67). An ANOVA revealed a significant main effect of Experiment (*F*(2, 87) = 3.32, *p* =0.041, *ηp2* = 0.071), with Bonferroni corrected post hoc comparisons indicating significant differences between Experiment 1 and Experiment 2 (*t*(2, 87) = -2.50, *p* = 0.043, *d* = 0.65, *CI* [0.62 26.83], *BF*_01_ = 0.500). Importantly, there were no significant differences between neither Experiments 1 and 3 (*t*(2, 87) = -1.80, *p* = 0.226, *d* = 0.47, *CI* [-3.21 22.99], *BF*_01_ = 0.744), nor Experiments 2 and 3 (*t*(2, 87) = -0.69, *p* = 1.000, *d* = -0.18, *CI* [-16.93 9.27], *BF*_01_ = 3.051) demonstrating that the attenuation observed in Experiment 3 cannot be attributed to time differences between the participants’ movement and the received touch.

**Supplemental Figure S1**

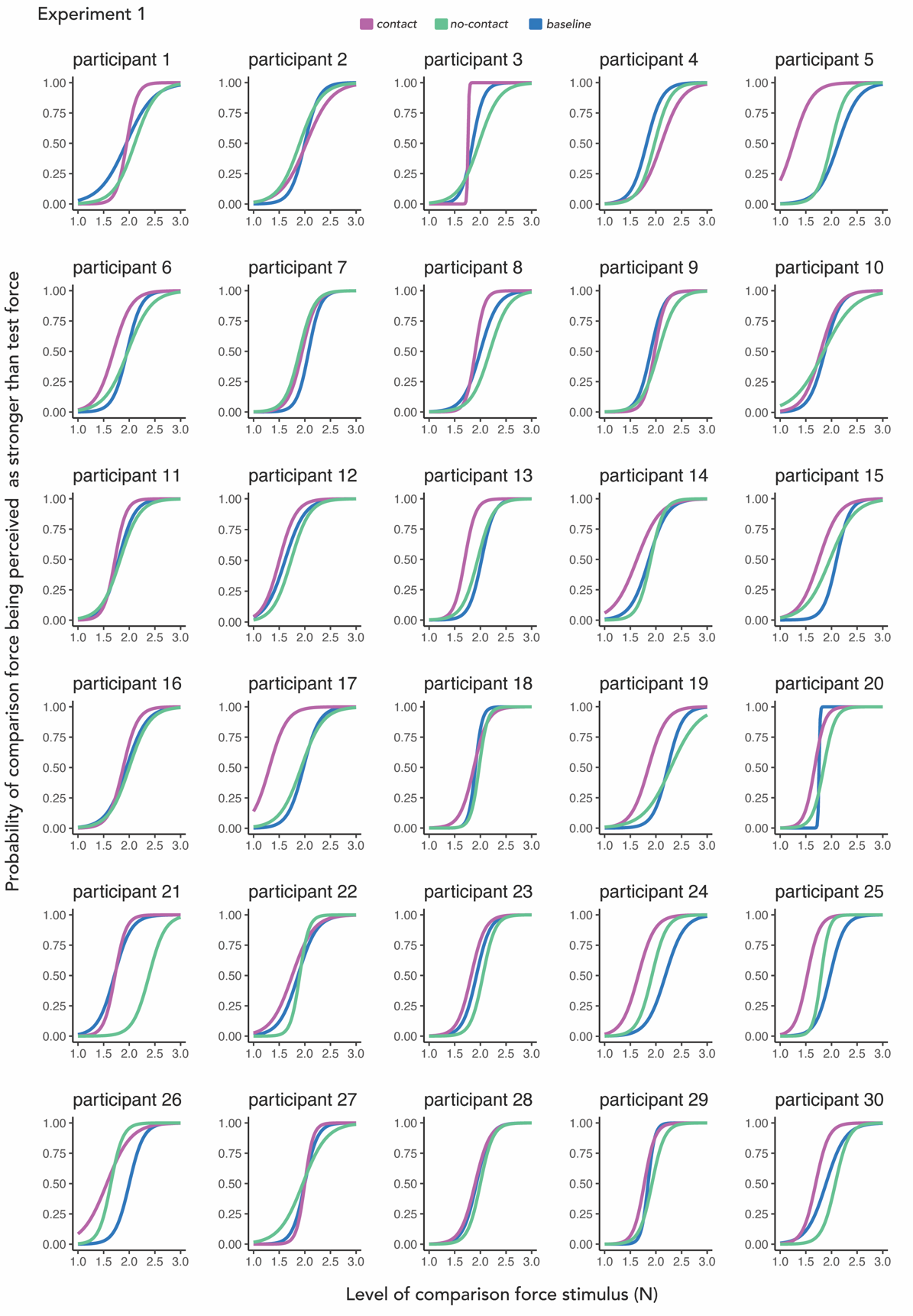

**Figure S1. Fitted logistic models based on the participants’ responses under each condition of Experiment 1.**

**Supplemental Figure S2**

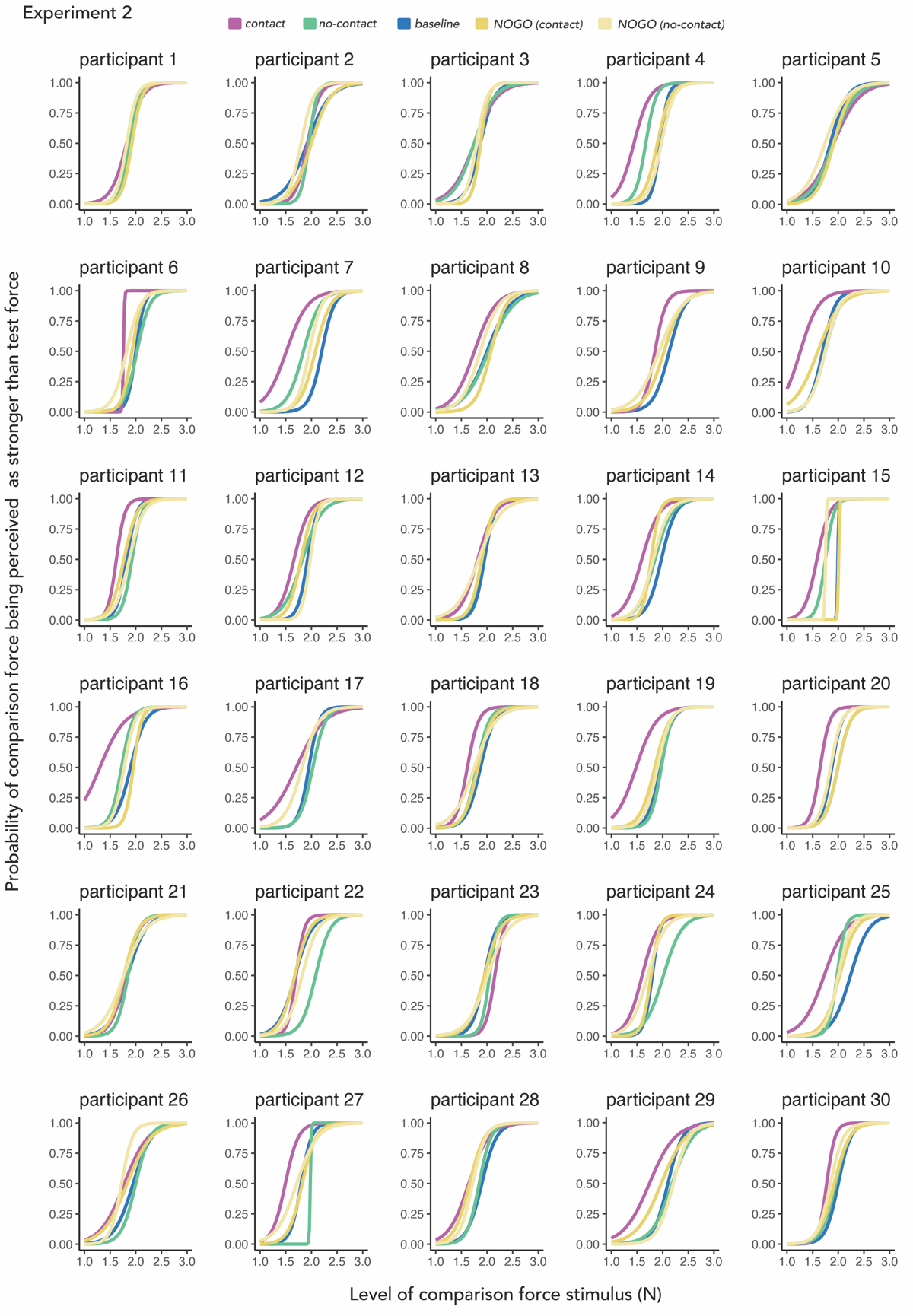

**Figure S2. Fitted logistic models based on the participants’ responses under each condition of Experiment 2.**

**Supplemental Figure S3**

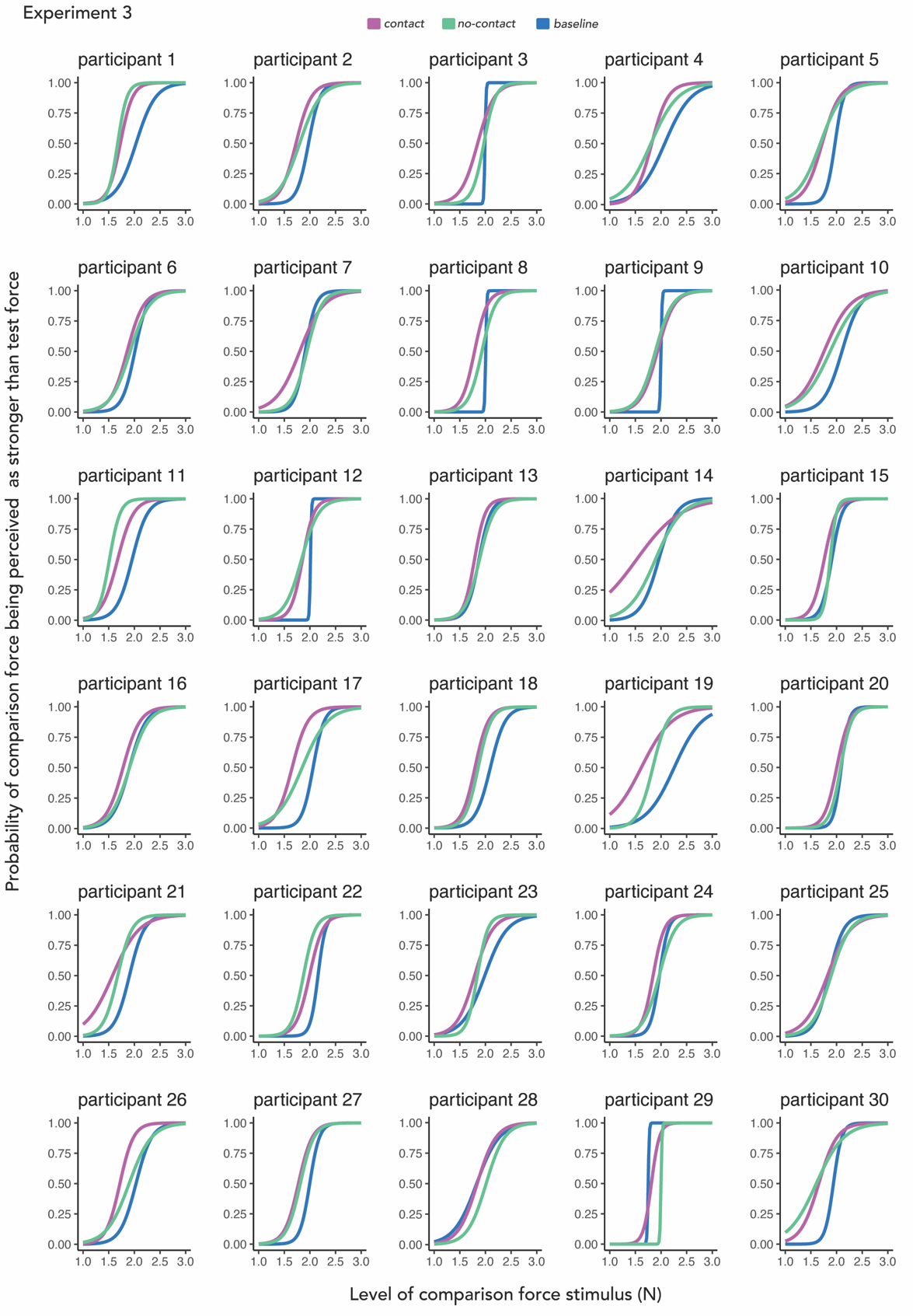

**Figure S3. Fitted logistic models based on the participants’ responses under each condition of Experiment 3.**
